# Magnetic-Alignment of Polymer Nanodiscs Probed by Solid-State NMR Spectroscopy

**DOI:** 10.1101/844332

**Authors:** Thirupathi Ravula, JaeWoong Kim, Dong-Kuk Lee, Ayyalusamy Ramamoorthy

## Abstract

The ability of amphipathic polymers to self-assemble with lipids and form nanodiscs has been a boon for the field of functional reconstitution of membrane proteins. In a field dominated by detergent micelles, a unique feature of polymer nanodiscs is their much-desired ability to align in the presence of an external magnetic field. Magnetic alignment facilitates the application of solid-state NMR spectroscopy and aids in the measurement of residual dipolar couplings (RDCs) via well-established solution NMR spectroscopy. In this study, we comprehensively investigate the magnetic-alignment properties of SMA-QA polymer based nanodiscs by using ^31^P and ^14^N solid-state NMR experiments under static conditions. The results reported herein demonstrate the spontaneous magnetic-alignment of large-size (≥ 20 nm diameter) SMA-QA nanodiscs (also called as *macro-nanodiscs*) with the lipid-bilayer-normal perpendicular to the magnetic field direction. Consequently, the orientation of *macro-nanodiscs* are further shown to flip their alignment axis parallel to the magnetic field direction upon the addition of a paramagnetic lanthanide salt. These results demonstrate the use of SMA-QA polymer nanodiscs for solid-state NMR applications including structural studies on membrane proteins.

## Introduction

Membrane proteins play a central and intricate part to many necessary cellular functions. However the study of membrane proteins is formidable due to a common loss of solubility, function, and structure when removed from a native-like membrane environment essential for membrane protein stability.^1–4^ To overcome this major hindrance in the membrane protein field, a major active area of research has been the development of new membrane mimetic systems.^1, 5–15^ While several solubilization systems have been introduced for the study membrane proteins (detergent micelles, bicelles, and nanodiscs),^5, 16^ nanodiscs have been proven to have advantages over other solubilization/membrane mimetics systems.^17–20^ Nanodiscs are lipid bilayer discs surrounded by amphiphilic macromolecules comprising of a protein,^18^ peptide,^21–23^ or polymer.^24^ Compared to other systems, polymer based nanodiscs have the unique capability of being able to extract membrane proteins directly from their native environment without the membrane proteins ever leaving the lipid bilayer.^25–26^ Recent developments in the polymer nanodiscs field expanded the applications of nanodiscs by using a wide variety of biophysical techniques to study membrane proteins.^17, 19, 27–39^

For accommodation of different sizes of membrane proteins and membrane associated protein-protein complexes, and to achieve magnetic-alignment, the size of nanodiscs should be variable. This is shown to be achievable with different polymer to lipid ratios.^40–41^ The large nanodiscs, called as *macro-nanodiscs*, (>20 nm diameter) that align in an external magnetic field^40, 42–43^ have been shown to be useful for structural studies using static solid-state NMR experiments^40, 44^ as well as an alignment medium to measure residual dipolar couplings via solution NMR experiments.^45^ In spite of such unique magnetic-alignment properties of nanodiscs, a comprehensive study on understanding the factors affecting the alignment of nanodiscs is lacking. In this study, we undertook a comprehensive list of experiments to study the magnetic-alignment of polymer based nanodiscs made from a positively charged polymer SMA-QA^43^ and DMPC lipids. Magnetic-alignment of SMA-QA+DMPC nanodiscs was studied using ^31^P and ^14^N solid-state NMR experiments. SMA-QA nanodiscs spontaneously align with the lipid bilayer normal perpendicular to the applied magnetic field direction. We also show that the nanodiscs alignment direction can be changed (or flipped) by the addition of lanthanide salts.

## Materials and Methods

Poly(Styrene-co-Maleic Anhydride) cumene terminated (SMA) with a ̴1.3:1 molar ratio of styrene:maleic anhydride and average molecular weight (M_n_) of 1600 g/mol, *N,N*-Dimethylformamide (DMF), Triethylamine (Et_3_N), HEPES, acetic acid (HOAc), hydrochloric acid (HCl), (2-Aminoethyl)trimethylammonium chloride hydrochloride, and sodium hydroxide (NaOH) were purchased from Sigma-Aldrich®. 1,2-dimyristoyl-*sn*-glycero-3-phosphocholine (DMPC) was purchased from Avanti Lipids Polar, Inc®.

### Synthesis and characterization of SMA-QA polymer

SMA-QA polymer was synthesized as reported previously.^43^ Briefly, 2 g of SMA and 2.86 g of (2-Aminoethyl)trimethylammonium chloride hydrochloride (15 eq) was dissolved in 50 mL of DMF, followed by the addition of 4 mL of Et_3_N (30 eq). The reaction mixture was stirred for 3 hrs at 80 °C. After 3 hrs the reaction mixture was cooled to room temperature and precipitated with ice-cooled diethyl ether. The resulting precipitate was washed three times with ether and dried under vacuum to give a white powder. The resulting powder was added to 2.25 g of sodium acetate, 50 mL of acetic anhydride and 6 mL of Et_3_N. The reaction mixture was stirred for 12 hrs at 80 °C. After the reaction, acetonitrile (50 mL) was added, discarded the white powder present at the bottom of the reaction mixture by carefully taking the supernatant with a pipette. The polymer was precipitated by the addition of diethyl ether. The resulting product was washed three times with ether and dried under vacuum to give a brown powder. The resulting crude polymer product was purified using LH-20 to remove the salts and lyophilized to produce 1.8 g brown powder of SMA-QA polymer.

### Preparation and characterization of polymer nanodiscs

In this study, we used two different approaches to prepare the nanodiscs samples. In the first approach, nanodiscs were formed by directly adding SMA-QA to DMPC vesicles (LUVs or MLVs) at a polymer:lipid ratio of 0.5:1 w/w and the resultant sample was directly used in experiments. In the second approach, the nanodisc sample system was prepared using the same first approach, however we included a size exclusion chromatography (SEC) purification step at the end to remove any non-nanodiscs forming free polymer from the solution as explained below (Figure 1). Both types of nanodiscs samples prepared were characterized dynamic light scattering (DLS) and transmission electron microscopy (TEM) experiments, and were investigated by solid-state NMR experiments under static conditions as explained in the results and discussion section below.

**Figure 1.**
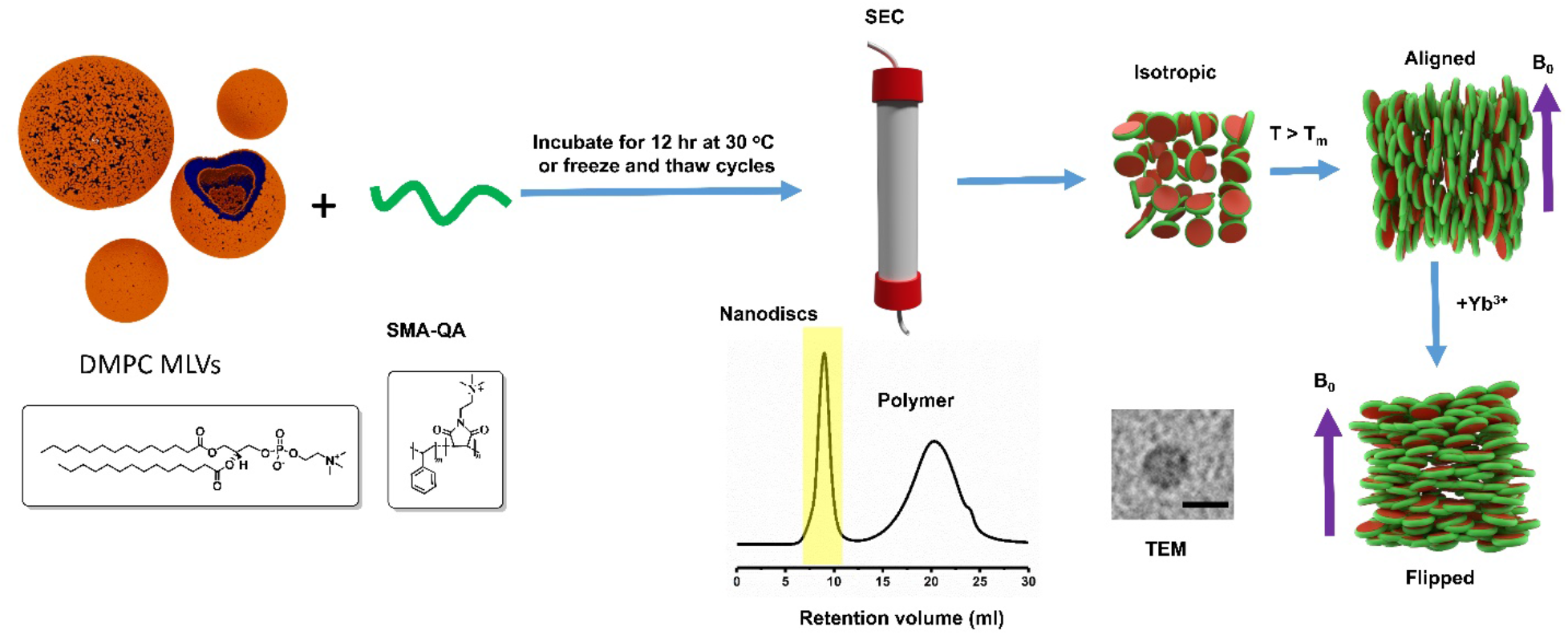
Nanodiscs preparation and characterization. (Top row) Schematic representation of the preparation of nanodiscs for solid-state NMR experiments, isotropic nanodiscs, and the alignment of macro-nanodiscs in the presence of an external magnetic field. (Bottom row) DMPC and SMA-QA chemical formulae, SEC chromatogram, TEM, and an illustration of flipped macro-nanodiscs in presence of lanthanide ions.

### NMR sample preparation

To prepare samples for solid-state NMR experiments, 8 mg of DMPC dissolved in CHCl_3_ was used for each sample preparation. The samples were dried under a stream of nitrogen gas, followed by overnight drying under vacuum (30 °C) to completely remove any residual solvent. Tris buffer (10 mM Tris, pH 7.4) was added to hydrate the lipid film, vortexed for 2 min above the lipid’s phase transition temperature and freeze-thawed using liquid nitrogen at least 4 times. To obtain uniform size of large unilamellar vesicles (LUVs) of 1 μm in diameter, the LUVs were extruded through polycarbonate filters (pore size of 1μm, Nuclepore®, Whatman, NJ, USA) mounted in a mini extruder (Avanti Polar Lipids, AL, USA) fitted with two 1.0 mL Hamilton gastight syringes (Hamilton, NV, USA). Samples were typically subjected to 23 passes through the filter. Odd number of passages were performed to avoid contamination of the sample by vesicles that have not passed through the first filter. SMA-QA sample, dissolved in Tris buffer, was added to LUVs to prepare the desired polymer concentration by making a final sample volume of 150 μL. The mixture was freeze-thawed using liquid nitrogen 3~5 times to get a transparent solution.

### ^31^P NMR experiments

Time dependent NMR experiments were performed on an Agilent NMR spectrometer operating at the resonance frequency of 699.88 MHz for ^1^ H and 283.31 MHz for ^31^P nuclei. A 4 mm triple-resonance HXY MAS NMR probe (Agilent) was used under static condition. ^31^P NMR spectra were acquired using a 5.5 μs 90° pulse followed by acquisition under 24 kHz TPPM proton decoupling. 128 scans were acquired for each sample with a relaxation/recycle delay time of 2.0 s. Each sample was put in a 4 mm pyrex glass tube, which was cut to fit into the 4 mm MAS probe. A Varian/Agilent temperature control unit was used to maintain the sample temperature. All ^31^P NMR spectra were processed using 150 Hz line broadening and referenced externally to 85% phosphoric acid (0 ppm). Temperature dependent experiments were performed on a Bruker NMR spectrometer at a resonance frequency of 400.11 MHz for proton and 161.97 MHz for ^31^P nuclei. 5 mm triple-resonance HXY MAS NMR probe was used under static conditions. ^31^P NMR spectra were acquired using a 5 μs 90° pulse followed by 25 kHz TPPM proton decoupling. 512 scans were acquired for each sample with a relaxation/recycle delay of 2.0 s.

### ^14^N NMR Experiments

Nitrogen-14 NMR spectra were acquired using a Bruker 400 MHz solid-state NMR spectrometer and a 5 mm double-resonance probe operating at the ^14^N resonance frequency of 28.910 MHz. ^14^N NMR spectra were recorded using the quadrupole-echo pulse sequence^46^ with a 90° pulse length of 8 μs and an echo-delay of 80 μs. ^14^N magnetization was acquired using 25 ms acquisition time, 20,000 scans and a recycle delay of 0.9 s with no ^1^H decoupling.

## Results and Discussion

In this study, we used nanodiscs composed of a positively charged SMA-QA polymer^43^ and DMPC lipids. SMA-QA was synthesized and characterized similar to the procedure as reported in our previous studies.^43^ Polymer macro-nanodiscs were prepared by the addition of the DMPC liposomes to the polymer stock solution of weight ratio 1:0.5 DMPC:SMA-QA to give the final lipid concentration of 100 mg/mL. The resulting solution was subjected to freeze thaw cycles until it became transparent, and the polymer nanodiscs solution was transferred to the NMR tube. To monitor the time-dependent changes in the alignment properties of the sample, a series of ^31^P NMR spectra were acquired as a function of time at 308 K (Figure 2). Initially, we observed a peak around ~-1 ppm and a small shoulder peak at ~-13 ppm. The observed peak at −1 ppm is from the isotropically tumbling nanodiscs, whereas the peak at −13 ppm is from the macro-nanodiscs aligned with the bilayer normal perpendicular to the magnetic field direction. The peak intensity for the aligned macro-nanodics increased whereas the isotropic peak intensity decreased as a function of time, and after 3 hrs the aligned peak became predominant. These results suggest that macro-nanodiscs possess a negative magnetic susceptibility and align in such a way that the lipid bilayer normal orients perpendicular to the magnetic field direction as shown in previous studies.^40, 43, 47^

**Figure 2.**
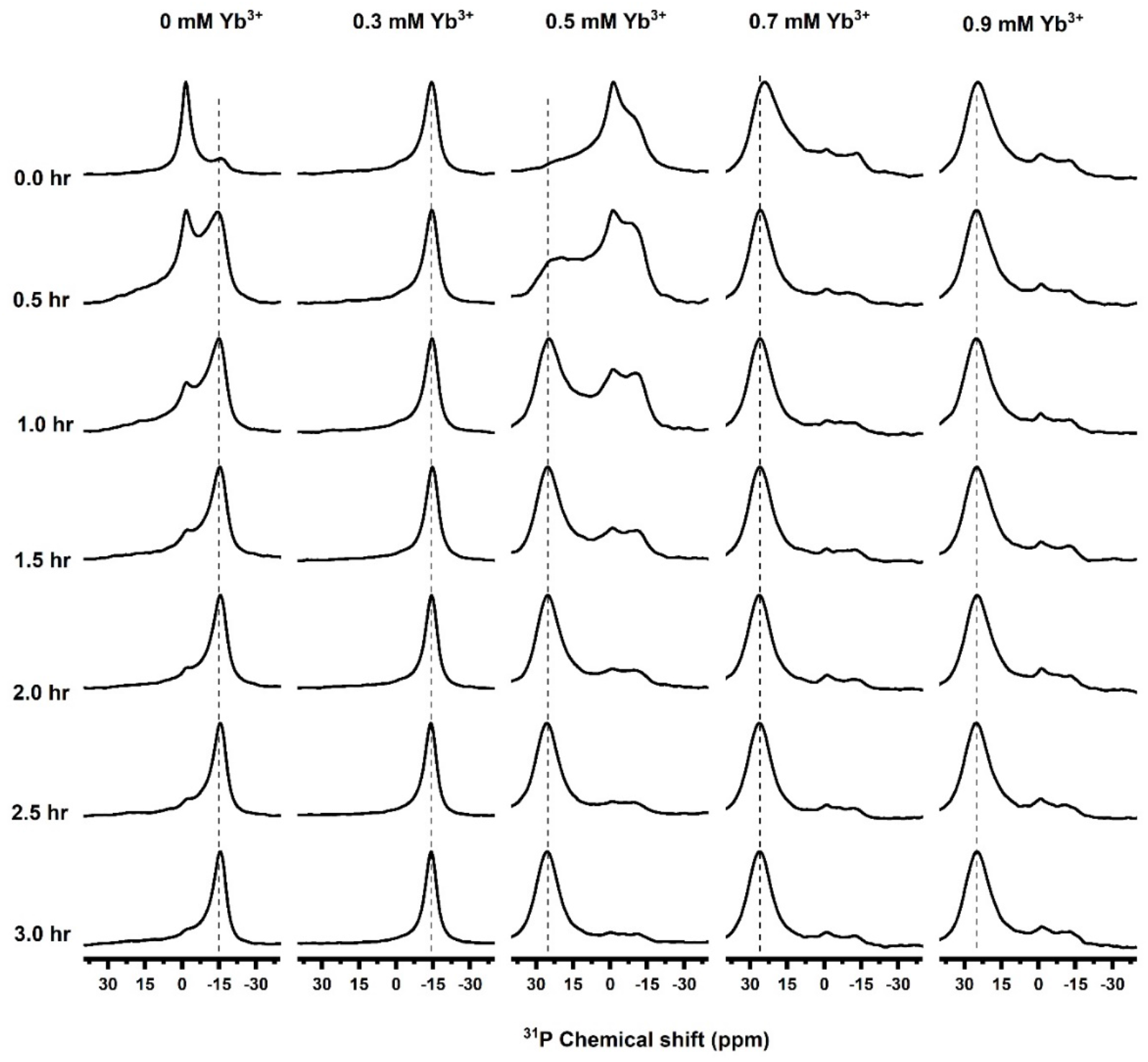
Magnetic alignment and flipping of macro-nanodiscs. ^31^P NMR spectra of unpurified SMA-QA:DMPC nanodiscs recorded as a function of time at 308 K, and at various concentrations of YbCl_3_. ^31^P peaks appearing at ~ −15, 0 and −22 ppm indicate a perpendicular orientation, isotropic phase, and a parallel orientation, respectively. Here, perpendicular and parallel orientations mean the orientation of the lipid bilayer normal relative to the direction of an external magnetic field. All of these NMR spectra were acquired using an Agilent 700 MHz NMR spectrometer.

The magnetic susceptibility of these nanodiscs can be changed by the addition of paramagnetic metal ions such as YbCl_3_ which has a positive magnetic susceptibility.^48–49^ To study the alignment behavior of macro-nanodiscs ^31^P NMR spectra were acquired as a function of time by adding different concentrations of YbCl_3_ to the aligned macro-nanodiscs. Upon the addition of sufficient Yb^3+^ ions, we observed a change in the orientation of macro-nanodiscs resulting in the bilayer normal parallel to the magnetic field direction. At a low concentration of YbCl_3_ (0.3 mM), no change in the ^31^P NMR spectrum was observed. But, as we increased the concentration of YbCl_3_ salt to 0.5 mM, a powder pattern along with an isotropic peak in the ^31^P NMR spectrum was observed initially. After 0.5 hrs a new peak appeared at 25 ppm whose intensity increased with further time, and it became the predominant peak after 3 hrs. This observation suggests that the macro-nanodiscs alignment is flipped to orient the bilayer normal parallel to the magnetic field direction. Further increase in the concentration of YbCl_3_ did not show any difference in the ^31^P peak intensity indicating no major changes in the alignment of macro-nanodiscs.

While the simple addition of polymer to lipids result in the formation of nanodiscs, there is always a portion of the added polymer that stays in solution. The free polymer in solution can be removed from the sample by using SEC to yield purified nanodiscs. We observed that the purified nanodiscs aligned much faster (< 1hr) as compared to the unpurified nanodiscs (figure 3). These nanodiscs exhibited a sharp peak at −1ppm suggesting isotropic nature of the macro-nanodiscs. As the temperature is increased above the gel-to-liquid crystalline phase transition temperature of the lipids (T > T_m_), an additional peak at −13 ppm in the ^31^P spectrum appeared suggesting the spontaneous magnetic-alignment of nanodiscs (Figure 3). As shown in Figure 3, at high temperatures (> 310 K), the isotropic peak at −1 ppm appeared with far less intensity suggesting significantly reduced isotropic population of macro-nanodiscs. In fact, for a range of temperatures (between 295 and 310 K), the ^31^P NMR spectra exhibited no isotropic behavior indicating an alignment of all macro-nanodiscs in the sample. ^31^P NMR spectra were also recorded upon the addition of different concentrations of YbCl_3_ as a function of temperature (Figure 3). For 0.5 and 1 mM concentrations of YbCl_3_, ^31^P NMR spectra showed a combination of powder pattern, isotropic peak, alignment with the bilayer normal perpendicular to the magnetic field direction, and flipping of nanodiscs with the bilayer normal parallel to the magnetic field direction depending on the temperature of the sample as shown in Figure 3. While the presence of 0.5 mM YbCl_3_ showed flipping of partial macro-nanodiscs population, the addition of a 1 mM YbCl_3_ showed complete flipping of nanodiscs at temperature 305 K. Further increase in temperature above 305 K showed the presence of an isotropic peak along with a peak at 21 ppm for samples containing YbCl_3_. At 1.5 mM YbCl_3_, the samples exhibited no significant powder pattern and aligned with the bilayer normal parallel to the magnetic field direction at 305 K. Thus, the ^31^P NMR experimental results presented in Figures 2 and 3 clearly demonstrate the magnetic-alignment of macro-nanodiscs and their direction of the alignment can be flipped by the addition of lanthanide ions like YbCl_3_. The observed chemical shifts are reported in Table 1. In addition, these results show that temperature and the concentration of lanthanide ions need to be optimized to achieve desirable magnetic-alignment of macro-nanodiscs. The sample and experimental conditions also depend on the type of lipids present in the nanodiscs. These behavior of macro-nanodiscs is similar to that of that of the well-studied bicelles samples.^47–48^

**Figure 3.**
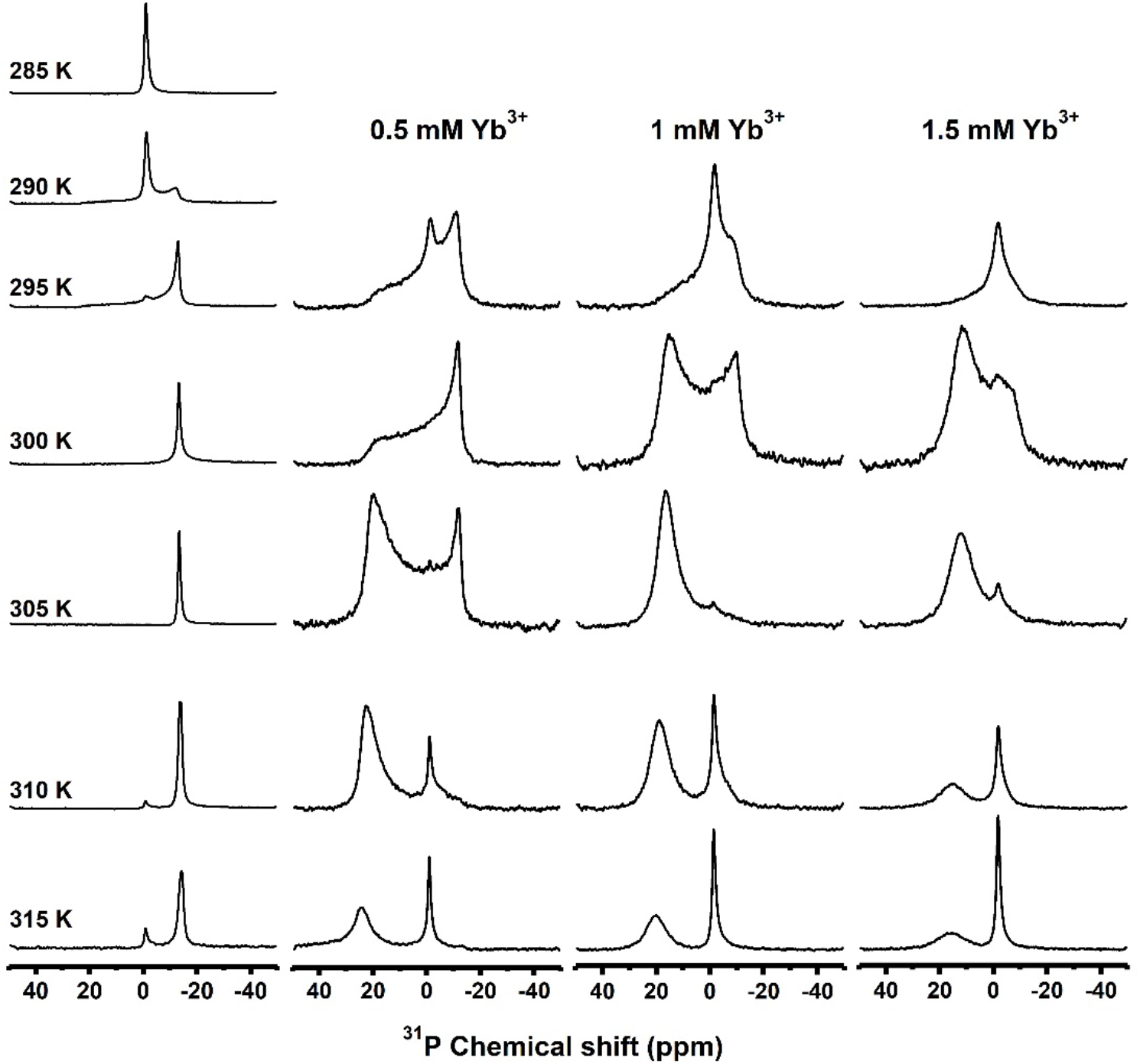
Temperature dependence of magnetic alignment and flipping of macro-nanodiscs. ^31^P NMR spectra of DMPC:SMA-QA (1:0.5 *w/w*) macro-nanodiscs in the absence (column 1) and presence of YbCl_3_ (columns 2-4) at different temperatures. A 100 mg/mL lipid concentration was used in the sample. All of these NMR spectra were acquired using a Bruker 400 MHz NMR spectrometer.

**Table 1.**
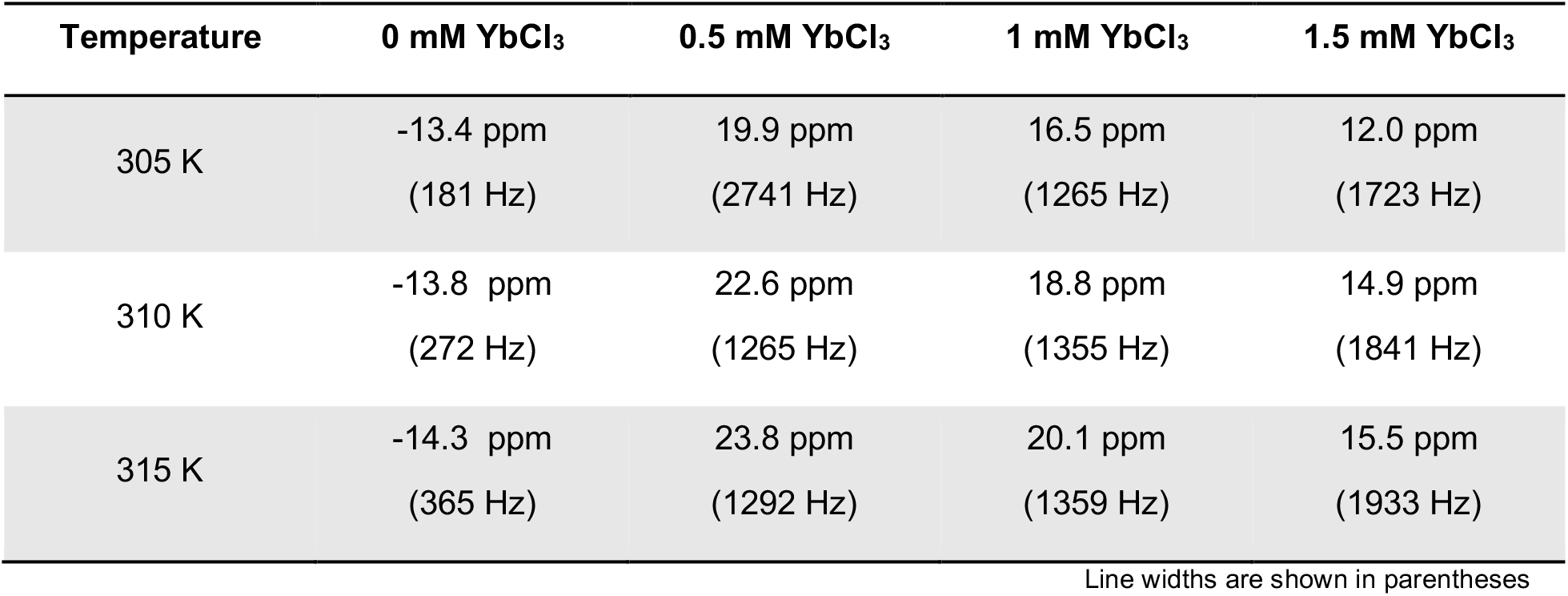
^31^P Chemical shifts and linewidths measured from NMR spectra of DMPC:SMA-QA (1:0.5 *w/w*) macro-nanodiscs in the absence (column 1) and presence of YbCl_3_ (columns 2-4) at different temperatures.

In addition to ^31^P NMR, ^14^N NMR spectroscopy is a powerful technique in sensing the membrane surface charge potential as well as useful to study the interaction of membrane active biomolecules due to the choline moiety.^50–55^ ^14^N is a quadrupolar nucleus with nuclear spin quantum number 1, and the quadrupole coupling constant depends on the orientation of the C-N bond vector of the choline group of DMPC with respect to the applied magnetic field direction, hydration, temperature, ions and other ligands, and mobility.^50–52^ ^14^N NMR experiments have previously been used to study the magnetic-alignment of bicelles.^51, 53–54^ ^14^N NMR spectra of macro-nanodiscs at 295 K showed the presence of a narrow peak centered at 0 kHz arising from the isotropic phase. As the temperature of the sample is increased to 310 K, the ^14^N NMR spectra showed the 0 kHz isotropic peak as well as the two additional peaks corresponding to a quadrupole splitting of 9.4 kHz. The quadrupolar splitting of 9.4 土 0.3 kHz arises from the magnetic-alignment of macro-nandiscs with the bilayer normal oriented perpendicular to the magnetic field axis. The small peak at 0 kHz may be assigned to the quaternary ammonium group from the SMA-QA polymer. With the addition of YbCl_3_, the choline group’s ^14^N quadrupolar splitting has doubled from 9.4 ± 0.3 kHz to 18.5 ± 0.5 kHz indicating the flip of nanodiscs alignment from perpendicular to parallel orientation of the bilayer normal with respect to the applied magnetic field axis. ^14^N NMR spectra acquired at different temperatures revealed macro-nanodiscs at low temperatures showed the presence of an isotropic phase whereas the increase in temperature (305 K and above) showed a quadrupolar splitting of ~18 kHz (Figure 4). Further increase in the sample temperature showed an increase in the intensity of the center peak at 0 kHz suggesting the coexistence of isotropic phase and flipped macro-nanodiscs. These results are in excellent agreement with the above presented ^31^P NMR results.

**Figure 4.**
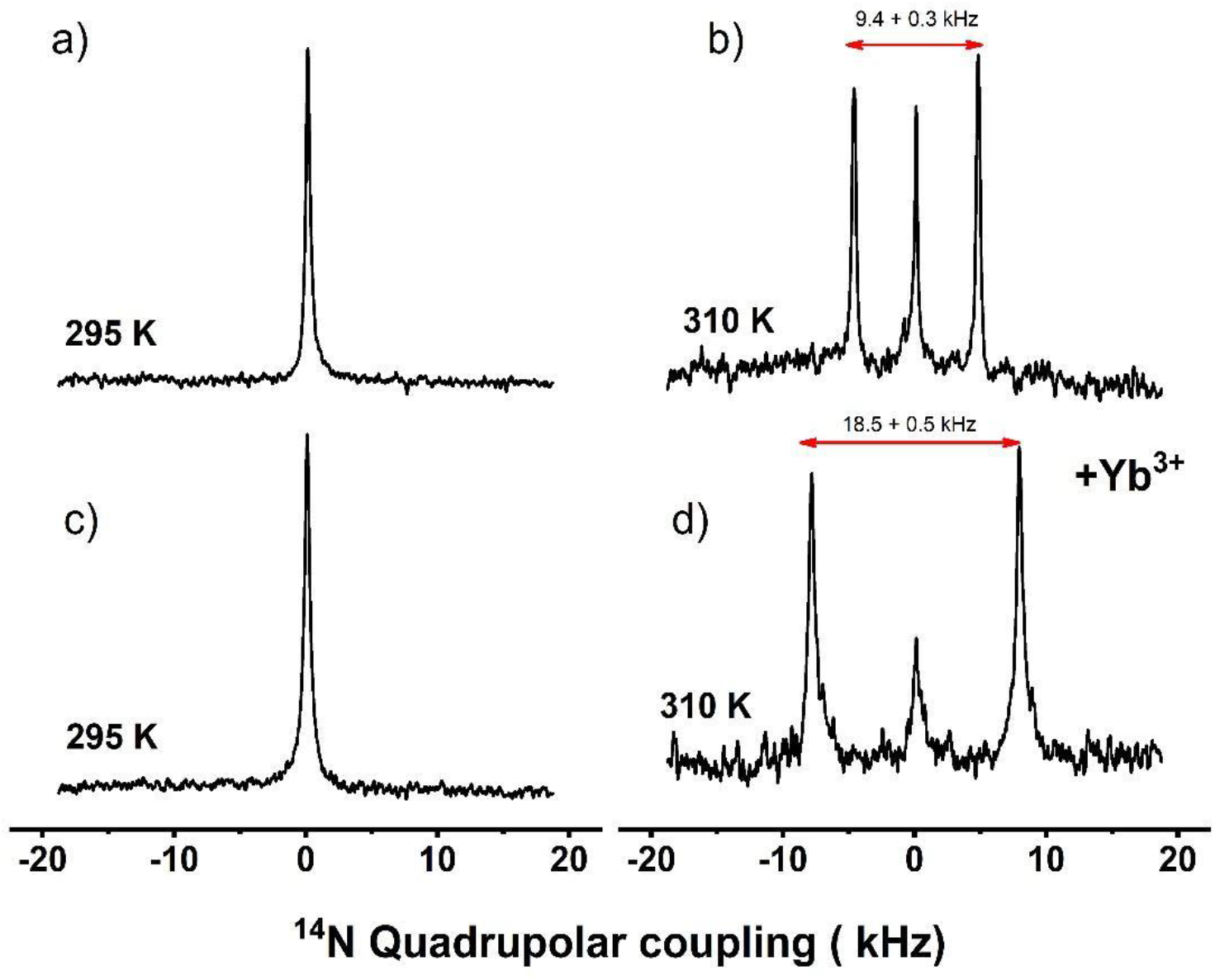
Magnetic alignment and flipping of macro-nanodiscs by ^14^N NMR. Nitrogen-14 NMR spectra of SMA-QA:DMPC with (c and d) and without (a and b) YbCl_3_ acquired at 295 K (a and c) and 310 K (b and d). The isotropic ^14^N NMR spectra (a and c) indicate the macro-nanodiscs are isotropic in the gel phase, i.e. below the gel to liquid crystalline phase transition temperature of DMPC lipids. The magnetic alignment (b) and its flipping due to the presence of 1 mM YbCl_3_ (d) are revealed by the observed ^14^N quadrupole splitting of 9.4 kHz (b) and 18.8 kHz (d) for the bilayer normal oriented perpendicular (b) and parallel (d) to the applied magnetic field direction, respectively. All of these NMR spectra were acquired using a Bruker 400 MHz NMR spectrometer.

**Figure 5.**
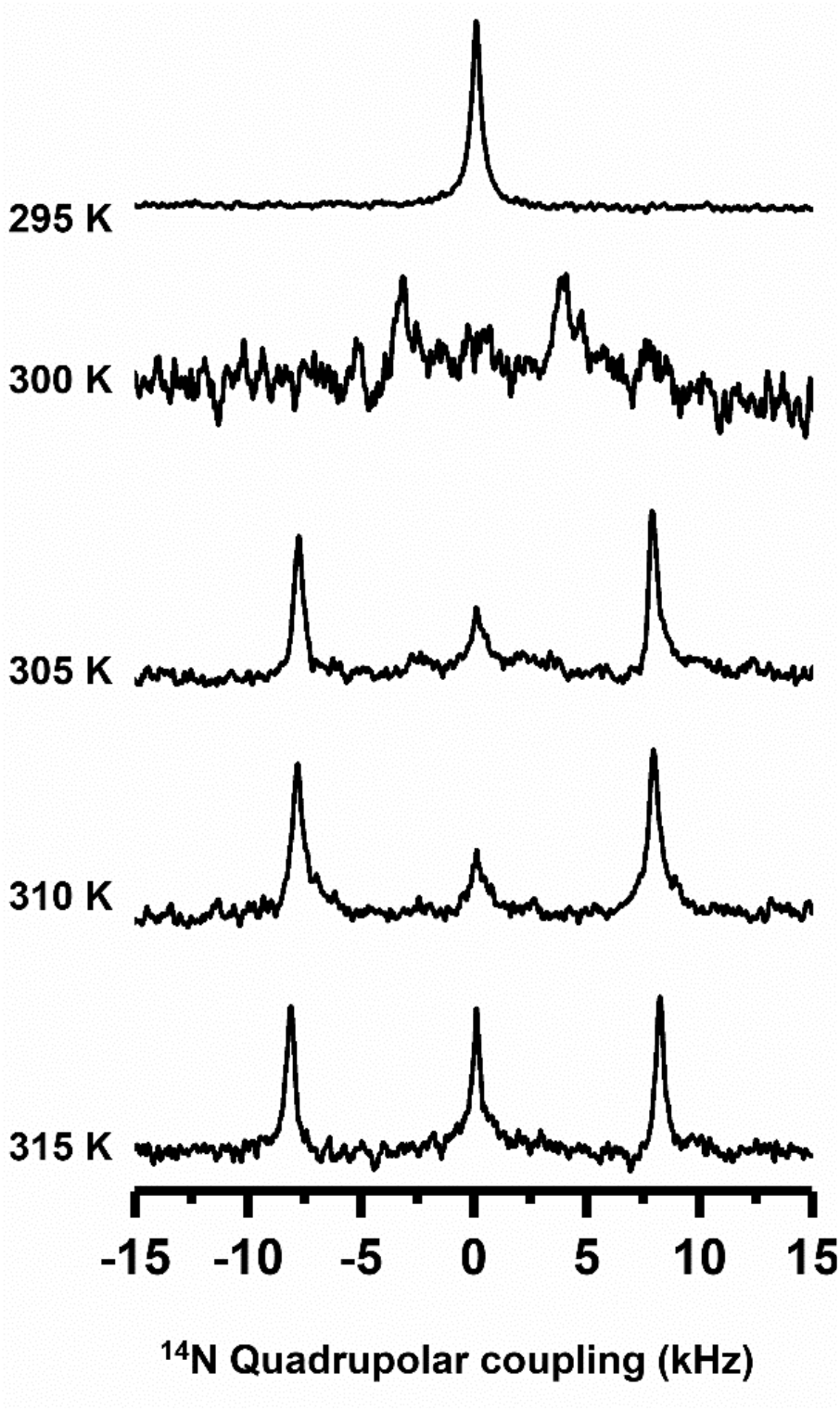
Temperature dependence of magnetic alignment of macro-nanodiscs by ^14^N NMR. Nitrogen-14 NMR spectra of flipped SMA-QA:DMPC macro-nanodiscs containing 1 mM YbCl_3_ as a function of temperature. Macro-nanodiscs are isotropic (and unaligned, indicated by 0 kHz ^14^N quadrupole coupling) below the gel to liquid crystalline phase transition temperature (295 K) of DMPC lipids. Above the phase transition temperature (>300 K), macro-nanodiscs align with the lipid bilayer normal parallel to the applied magnetic field direction in presence of 1 mM YbCl_3_ as revealed by the observed quadrupole splitting of 18.8 kHz. All of these NMR spectra were acquired using a Bruker 400 MHz NMR spectrometer.

## Conclusion

Polymer nanodiscs are a valuable tool used to study membrane proteins. With the ability to align in the presence of a magnetic field such that the bilayer normal is oriented perpendicular to the magnetic field direction, macro-nanodiscs show an promising use in solid-state NMR studies.^14^N NMR spectra reported in this study reveal the spontaneous alignment and the time and temperature required to align in the presence of an external magnetic field. Our results successfully demonstrate the ability of YbCl_3_ ions to flip the direction of macro-nanodiscs. We believe that this study will be useful in optimizing the conditions that are needed to study membrane proteins reconstituted in polymer nanodiscs by solid-state NMR spectroscopy.^56–60^ It is also worth mentioning that pH tolerance and deviant metal-ions resistance of SMA-QA polymer nanodiscs further expands the applications for solid-state NMR applications including structural studies on membrane proteins. In addition, as shown in our recent study, magnetically-aligned polymer macro-nanodiscs can be used to measure residual-dipolar-couplings by well-established solution NMR methods to study the structure and dynamics of water-soluble molecules including proteins, peptides, DNA, RNA and small molecule compounds.^61–63^ We expect these unique properties of polymer based macro-nanodiscs to open avenues to expand the applications of both solution and solid-state NMR techniques, and create opportunities for further development of novel NMR approaches.^2, 64–67^

## Acknowledgements

This study was supported by NIH (GM084018 and AG048934 to A.R.). We thank Nathaniel Hardin for his help in preparing the manuscript.

## References

1. Garavito, R. M.; Ferguson-Miller, S., Detergents as tools in membrane biochemistry. J Biol Chem 2001, 276, 32403–6.

2. Xiao, P.; Bolton, D.; Munro, R. A.; Brown, L. S.; Ladizhansky, V., Solid-state NMR spectroscopy based atomistic view of a membrane protein unfolding pathway. Nat Commun 2019, 10, 3867.

3. Wylie, B. J.; Bhate, M. P.; McDermott, A. E., Transmembrane allosteric coupling of the gates in a potassium channel. Proc Natl Acad Sci U S A 2014, 111, 185–90.

4. Chipot, C.; Dehez, F.; Schnell, J. R.; Zitzmann, N.; Pebay-Peyroula, E.; Catoire, L. J.; Miroux, B.; Kunji, E. R. S.; Veglia, G.; Cross, T. A.; Schanda, P., Perturbations of Native Membrane Protein Structure in Alkyl Phosphocholine Detergents: A Critical Assessment of NMR and Biophysical Studies. Chemical Reviews 2018, 118, 3559–3607.

5. Frey, L.; Lakomek, N. A.; Riek, R.; Bibow, S., Micelles, Bicelles, and Nanodiscs: Comparing the Impact of Membrane Mimetics on Membrane Protein Backbone Dynamics. Angew Chem Int Ed Engl 2017, 56, 380–383.

6. Lee, S. C.; Pollock, N. L., Membrane proteins: is the future disc shaped? Biochem Soc Trans 2016, 44, 1011–8.

7. Helenius, A.; Simons, K., Solubilization of membranes by detergents. Biochim Biophys Acta 1975, 415, 29–79.

8. Seddon, A. M.; Curnow, P.; Booth, P. J., Membrane proteins, lipids and detergents: not just a soap opera. Biochim Biophys Acta 2004, 1666, 105–17.

9. Tanford, C.; Reynolds, J. A., Characterization of membrane proteins in detergent solutions. Biochim Biophys Acta 1976, 457, 133–70.

10. Czerski, L.; Sanders, C. R., Functionality of a membrane protein in bicelles. Anal Biochem 2000, 284, 327–33.

11. Parmar, M. J.; Lousa Cde, M.; Muench, S. P.; Goldman, A.; Postis, V. L., Artificial membranes for membrane protein purification, functionality and structure studies. Biochem Soc Trans 2016, 44, 877–82.

12. Kalipatnapu, S.; Chattopadhyay, A., Membrane protein solubilization: recent advances and challenges in solubilization of serotonin1A receptors. IUBMB Life 2005, 57, 505–12.

13. Rigaud, J.-L.; Lévy, D., Reconstitution of Membrane Proteins into Liposomes. In Methods Enzymol., Academic Press: 2003; Vol. 372, pp 65–86.

14. Tribet, C.; Audebert, R.; Popot, J. L., Amphipols: Polymers that keep membrane proteins soluble in aqueous solutions. Proc Natl Acad Sci U S A 1996, 93, 15047–15050.

15. Sanders, C. R.; Prosser, R. S., Bicelles: a model membrane system for all seasons? Structure 1998, 6, 1227–1234.

16. Morrison, E. A.; DeKoster, G. T.; Dutta, S.; Vafabakhsh, R.; Clarkson, M. W.; Bahl, A.; Kern, D.; Ha, T.; Henzler-Wildman, K. A., Antiparallel EmrE exports drugs by exchanging between asymmetric structures. Nature 2011, 481, 45–50.

17. Denisov, I. G.; Sligar, S. G., Nanodiscs in Membrane Biochemistry and Biophysics. Chem Rev 2017, 117, 4669–4713.

18. Denisov, I. G.; Grinkova, Y. V.; Lazarides, A. A.; Sligar, S. G., Directed self-assembly of monodisperse phospholipid bilayer Nanodiscs with controlled size. J Am Chem Soc 2004, 126, 3477–87.

19. Denisov, I. G.; Sligar, S. G., Nanodiscs for structural and functional studies of membrane proteins. Nat Struct Mol Biol 2016, 23, 481–6.

20. Nath, A.; Atkins, W. M.; Sligar, S. G., Applications of phospholipid bilayer nanodiscs in the study of membranes and membrane proteins. Biochemistry 2007, 46, 2059–69.

21. Kariyazono, H.; Nadai, R.; Miyajima, R.; Takechi-Haraya, Y.; Baba, T.; Shigenaga, A.; Okuhira, K.; Otaka, A.; Saito, H., Formation of stable nanodiscs by bihelical apolipoprotein A-I mimetic peptide. J Pept Sci 2016, 22, 116–22.

22. Mishra, V. K.; Palgunachari, M. N.; Krishna, R.; Glushka, J.; Segrest, J. P.; Anantharamaiah, G. M., Effect of leucine to phenylalanine substitution on the nonpolar face of a class A amphipathic helical peptide on its interaction with lipid: high resolution solution NMR studies of 4F-dimyristoylphosphatidylcholine discoidal complex. J Biol Chem 2008, 283, 34393–402.

23. Zhang, M.; Huang, R.; Ackermann, R.; Im, S. C.; Waskell, L.; Schwendeman, A.; Ramamoorthy, A., Reconstitution of the Cytb5-CytP450 Complex in Nanodiscs for Structural Studies Using NMR Spectroscopy. Angew. Chem. Int. Ed. 2016, 128, 4497–4499.

24. Knowles, T. J.; Finka, R.; Smith, C.; Lin, Y. P.; Dafforn, T.; Overduin, M., Membrane proteins solubilized intact in lipid containing nanoparticles bounded by styrene maleic acid copolymer. J Am Chem Soc 2009, 131, 7484–5.

25. Lee, S. C.; Knowles, T. J.; Postis, V. L.; Jamshad, M.; Parslow, R. A.; Lin, Y. P.; Goldman, A.; Sridhar, P.; Overduin, M.; Muench, S. P.; Dafforn, T. R., A method for detergent-free isolation of membrane proteins in their local lipid environment. Nat Protoc 2016, 11, 1149–62.

26. Dorr, J. M.; Koorengevel, M. C.; Schafer, M.; Prokofyev, A. V.; Scheidelaar, S.; van der Cruijsen, E. A.; Dafforn, T. R.; Baldus, M.; Killian, J. A., Detergent-free isolation, characterization, and functional reconstitution of a tetrameric K+ channel: the power of native nanodiscs. Proc Natl Acad Sci U S A 2014, 111, 18607–12.

27. Hagn, F.; Etzkorn, M.; Raschle, T.; Wagner, G., Optimized phospholipid bilayer nanodiscs facilitate high-resolution structure determination of membrane proteins. J Am Chem Soc 2013, 135, 1919–25.

28. Nasr, M. L.; Wagner, G., Covalently circularized nanodiscs; challenges and applications. Curr Opin Struct Biol 2018, 51, 129–134.

29. Denisov, I. G.; Sligar, S. G., Nanodiscs for structural and functional studies of membrane proteins. Nat. Struct. Mol. Biol. 2016, 23, 481.

30. Redhair, M.; Clouser, A. F.; Atkins, W. M., Hydrogen-deuterium exchange mass spectrometry of membrane proteins in lipid nanodiscs. Chem Phys Lipids 2019, 220, 14–22.

31. Overduin, M.; Esmaili, M., Structures and Interactions of Transmembrane Targets in Native Nanodiscs. SLAS Discov 2019, DOI:10.1177/2472555219857691.

32. Sun, C.; Benlekbir, S.; Venkatakrishnan, P.; Wang, Y.; Hong, S.; Hosler, J.; Tajkhorshid, E.; Rubinstein, J. L.; Gennis, R. B., Structure of the alternative complex III in a supercomplex with cytochrome oxidase. Nature 2018, 557, 123–126.

33. Orwick, M. C.; Judge, P. J.; Procek, J.; Lindholm, L.; Graziadei, A.; Engel, A.; Grobner, G.; Watts, A., Detergent-free formation and physicochemical characterization of nanosized lipid-polymer complexes: Lipodisq. Angew Chem Int Ed Engl 2012, 51, 4653–7.

34. Kopf, A. H.; Koorengevel, M. C.; van Walree, C. A.; Dafforn, T. R.; Killian, J. A., A simple and convenient method for the hydrolysis of styrene-maleic anhydride copolymers to styrene-maleic acid copolymers. Chem Phys Lipids 2019, 218, 85–90.

35. Dominguez Pardo, J. J.; van Walree, C. A.; Egmond, M. R.; Koorengevel, M. C.; Killian, J. A., Nanodiscs bounded by styrene-maleic acid allow trans-cis isomerization of enclosed photoswitches of azobenzene labeled lipids. Chem Phys Lipids 2019, 220, 1–5.

36. Danielczak, B.; Meister, A.; Keller, S., Influence of Mg(2+) and Ca(2+) on nanodisc formation by diisobutylene/maleic acid (DIBMA) copolymer. Chem Phys Lipids 2019, 221, 30–38.

37. Danielczak, B.; Keller, S., Collisional lipid exchange among DIBMA-encapsulated nanodiscs (DIBMALPs). European Polymer Journal 2018, 109, 206–213.

38. Oluwole, A. O.; Klingler, J.; Danielczak, B.; Babalola, J. O.; Vargas, C.; Pabst, G.; Keller, S., Formation of Lipid-Bilayer Nanodiscs by Diisobutylene/Maleic Acid (DIBMA) Copolymer. Langmuir 2017, 33, 14378–14388.

39. Oluwole, A. O.; Danielczak, B.; Meister, A.; Babalola, J. O.; Vargas, C.; Keller, S., Solubilization of Membrane Proteins into Functional Lipid-Bilayer Nanodiscs Using a Diisobutylene/Maleic Acid Copolymer. Angew Chem Int Ed Engl 2017, 56, 1919–1924.

40. Ravula, T.; Ramadugu, S. K.; Di Mauro, G.; Ramamoorthy, A., Bioinspired, Size-Tunable Self-Assembly of Polymer-Lipid Bilayer Nanodiscs. Angew Chem Int Ed Engl 2017, 56, 11466–11470.

41. Zhang, R.; Sahu, I. D.; Liu, L.; Osatuke, A.; Comer, R. G.; Dabney-Smith, C.; Lorigan, G. A., Characterizing the structure of lipodisq nanoparticles for membrane protein spectroscopic studies. Biochim Biophys Acta 2015, 1848, 329–33.

42. Radoicic, J.; Park, S. H.; Opella, S. J., Macrodiscs Comprising SMALPs for Oriented Sample Solid-State NMR Spectroscopy of Membrane Proteins. Biophys J 2018, 115, 22–25.

43. Ravula, T.; Hardin, N. Z.; Ramadugu, S. K.; Cox, S. J.; Ramamoorthy, A., Formation of pH-Resistant Monodispersed Polymer-Lipid Nanodiscs. Angew Chem Int Ed Engl 2018, 57, 1342–1345.

44. Ravula, T.; Hardin, N. Z.; Ramamoorthy, A., Polymer nanodiscs: Advantages and limitations. Chem Phys Lipids 2019, 219, 45–49.

45. Ravula, T.; Ramamoorthy, A., Magnetic Alignment of Polymer Macro-Nanodiscs Enables Residual-Dipolar-Coupling-Based High-Resolution Structural Studies by NMR Spectroscopy. Angew Chem Int Ed Engl 2019, 58, 14925–14928.

46. Weisman, I. D.; Bennett, L. H., Quadrupolar Echoes in Solids. Physical Review 1969, 181, 1341–1350.

47. Sanders, C. R.; Hare, B. J.; Howard, K. P.; Prestegard, J. H., Magnetically-Oriented Phospholipid Micelles as a Tool for the Study of Membrane-Associated Molecules. Prog Nucl Mag Res Sp 1994, 26, 421–444.

48. Prosser, R. S.; Hwang, J. S.; Vold, R. R., Magnetically aligned phospholipid bilayers with positive ordering: a new model membrane system. Biophys J 1998, 74, 2405–18.

49. Prosser, R. S.; Hunt, S. A.; DiNatale, J. A.; Vold, R. R., Magnetically Aligned Membrane Model Systems with Positive Order Parameter: Switching the Sign ofSzzwith Paramagnetic Ions. J Am Chem Soc 1996, 118, 269–270.

50. Lindstrom, F.; Williamson, P. T.; Grobner, G., Molecular insight into the electrostatic membrane surface potential by 14n/31p MAS NMR spectroscopy: nociceptin-lipid association. J Am Chem Soc 2005, 127, 6610–6.

51. Ramamoorthy, A.; Lee, D. K.; Santos, J. S.; Henzler-Wildman, K. A., Nitrogen-14 solid-state NMR spectroscopy of aligned phospholipid bilayers to probe peptide-lipid interaction and oligomerization of membrane associated peptides. J Am Chem Soc 2008, 130, 11023–9.

52. Ramamoorthy, A.; Thennarasu, S.; Lee, D. K.; Tan, A.; Maloy, L., Solid-state NMR investigation of the membrane-disrupting mechanism of antimicrobial peptides MSI-78 and MSI-594 derived from magainin 2 and melittin. Biophys J 2006, 91, 206–16.

53. Smith, P. E.; Brender, J. R.; Ramamoorthy, A., Induction of negative curvature as a mechanism of cell toxicity by amyloidogenic peptides: the case of islet amyloid polypeptide. J Am Chem Soc 2009, 131, 4470–8.

54. Semchyschyn, D. J.; Macdonald, P. M., Conformational response of the phosphatidylcholine headgroup to bilayer surface charge: torsion angle constraints from dipolar and quadrupolar couplings in bicelles. Magn Reson Chem 2004, 42, 89–104.

55. Scherer, P. G.; Seelig, J., Electric charge effects on phospholipid headgroups. Phosphatidylcholine in mixtures with cationic and anionic amphiphiles. Biochemistry 1989, 28, 7720–8.

56. Aisenbrey, C.; Salnikov, E. S.; Bechinger, B., Solid-State NMR Investigations of the MHC II Transmembrane Domains: Topological Equilibria and Lipid Interactions. J Membr Biol 2019, 252, 371–384.

57. Salnikov, E. S.; De Zotti, M.; Bobone, S.; Mazzuca, C.; Raya, J.; Siano, A. S.; Peggion, C.; Toniolo, C.; Stella, L.; Bechinger, B., Trichogin GA IV Alignment and Oligomerization in Phospholipid Bilayers. Chembiochem 2019, 20, 2141–2150.

58. Grage, S. L.; Sani, M. A.; Cheneval, O.; Henriques, S. T.; Schalck, C.; Heinzmann, R.; Mylne, J. S.; Mykhailiuk, P. K.; Afonin, S.; Komarov, I. V.; Separovic, F.; Craik, D. J.; Ulrich, A. S., Orientation and Location of the Cyclotide Kalata B1 in Lipid Bilayers Revealed by Solid-State NMR. Biophys J 2017, 112, 630–642.

59. Morton, C. J.; Sani, M. A.; Parker, M. W.; Separovic, F., Cholesterol-Dependent Cytolysins: Membrane and Protein Structural Requirements for Pore Formation. Chem Rev 2019, 119, 7721–7736.

60. Laadhari, M.; Arnold, A. A.; Gravel, A. E.; Separovic, F.; Marcotte, I., Interaction of the antimicrobial peptides caerin 1.1 and aurein 1.2 with intact bacteria by (2)H solid-state NMR. Biochim Biophys Acta 2016, 1858, 2959–2964.

61. Tjandra, N.; Bax, A., Direct Measurement of Distances and Angles in Biomolecules by NMR in a Dilute Liquid Crystalline Medium. Science 1997, 278, 1111.

62. Peti, W.; Meiler, J.; Bruschweiler, R.; Griesinger, C., Model-free analysis of protein backbone motion from residual dipolar couplings. J Am Chem Soc 2002, 124, 5822–33.

63. Liu, Y.; Navarro-Vazquez, A.; Gil, R. R.; Griesinger, C.; Martin, G. E.; Williamson, R. T., Application of anisotropic NMR parameters to the confirmation of molecular structure. Nat Protoc 2019, 14, 217–247.

64. Wang, S.; Gopinath, T.; Veglia, G., Improving the quality of oriented membrane protein spectra using heat-compensated separated local field experiments. J Biomol NMR 2019, DOI:10.1007/s10858-019-00273-1.

65. Hong, M.; Su, Y., Structure and dynamics of cationic membrane peptides and proteins: insights from solid-state NMR. Protein Sci 2011, 20, 641–55.

66. Leninger, M.; Sae Her, A.; Traaseth, N. J., Inducing conformational preference of the membrane protein transporter EmrE through conservative mutations. Elife 2019, 8, 48909.

67. Pinto, C.; Mance, D.; Sinnige, T.; Daniels, M.; Weingarth, M.; Baldus, M., Formation of the beta-barrel assembly machinery complex in lipid bilayers as seen by solid-state NMR. Nat Commun 2018, 9, 4135.

